# Pilot production of SARS-CoV-2 related proteins in plants: a proof of concept for rapid repurposing of indoors farms into biomanufacturing facilities

**DOI:** 10.1101/2020.10.13.331306

**Authors:** Borja Diego-Martin, Beatriz González, Marta Vazquez-Vilar, Sara Selma, Rubén Mateos-Fernández, Silvia Gianoglio, Asun Fernández-del-Carmen, Diego Orzáez

## Abstract

The current CoVid-19 crisis is revealing the strengths and the weaknesses of the world’s capacity to respond to a global health crisis. A critical weakness has resulted from the excessive centralization of the current biomanufacturing capacities, a matter of great concern, if not a source of nationalistic tensions. On the positive side, scientific data and information have been shared at an unprecedented speed fuelled by the preprint phenomena, and this has considerably strengthened our ability to develop new technology-based solutions. In this work we explore how, in a context of rapid exchange of scientific information, plant biofactories can serve as a rapid and easily adaptable solution for local manufacturing of bioreagents, more specifically recombinant antibodies. For this purpose, we tested our ability to produce, in the framework of an academic lab and in a matter of weeks, milligram amounts of six different recombinant monoclonal antibodies against SARS-CoV-2 in *Nicotiana benthamiana.* For the design of the antibodies we took advantage, among other data sources, of the DNA sequence information made rapidly available by other groups in preprint publications. mAbs were all engineered as single-chain fragments fused to a human gamma Fc and transiently expressed using a viral vector. In parallel, we also produced the recombinant SARS-CoV-2 N protein and its Receptor Binding Domain (RBD) *in planta* and used them to test the binding specificity of the recombinant mAbs. Finally, for two of the antibodies we assayed a simple scale-up production protocol based on the extraction of apoplastic fluid. Our results indicate that gram amounts of anti-SARS-CoV-2 antibodies could be easily produced in little more than 6 weeks in repurposed greenhouses with little infrastructure requirements using *N. benthamiana* as production platform. Similar procedures could be easily deployed to produce diagnostic reagents and, eventually, could be adapted for rapid therapeutic responses.

## INTRODUCTION

The current pandemic is evidencing several weaknesses in our ability to respond to a global crisis, one of which is the insufficient and heavily centralized distribution of the world manufacturing capacity of bioproducts such as antibodies, vaccines and other biological reagents, specially proteins. Since it is economically impracticable to ensure readiness by maintaining inactive infrastructures during large periods of normality, the development of dual-use systems has been proposed, which would serve regular production needs in normal times but could be rapidly repurposed to strategic manufacturing requirements in times of crisis. Ideally, such adaptable infrastructures should be widespread to serve local demand in case of emergency.

Recombinant protein production in plants is a technologically mature bioengineering discipline, with most current plant-based bioproduction platforms making use of non-food crops, mainly the *Nicotiana* species tabacum and *N. benthamiana* as biomanufacturing chassis (Moon et al., 2019; Capell et al., 2020). *N. benthamiana* is most frequently used in association with *Agrobacterium-mediated* transient expression, also known as agroinfiltration, a technology that dramatically reduces the time required for product development. Briefly, agroinfiltration consists in the massive delivery of an *Agrobacterium* suspension culture to the intercellular space of plant leaves, either by pressure, using a syringe (small scale), or applying vacuum to plants whose aerial parts have been submerged in a diluted *Agrobacterium* culture (large scale). *Agrobacterium* transfers its T-DNA to the cell nucleus, therefore massively reprogramming the plant cell machinery towards the synthesis of the T-DNA-encoded protein(s)-of-interest. Transient expression of the transgene is often assisted by self-replicating deconstructed virus vectors that amplify the transgene dose, thus boosting protein production by several orders of magnitude (Gleba et al., 2004). Other systems, such as the pEAQ system, rely on viral genetic elements for boosting expression without recurring to viral replication (Sainsbury et al., 2009). Transient expression in *N. benthamiana* has become the standard in plant-based recombinant protein production due to a unique combination of advantages, with speed and high yield as the most obvious ones. Maximum production levels in the g/Kg fresh weight (FW) range for certain highly stable proteins such as antibodies have been reported (Marillonnet et al., 2005). Regarding speed, the *in-planta* incubation times required to obtain maximum yield of recombinant protein are no more than two weeks.

An important, often insufficiently highlighted feature of *N. benthamiana* transient expression is its relatively small infrastructure requirements, partially overlapping with those employed in more conventional, medium/high-tech indoors agriculture, such as hydroponics, vertical farming, etc. (Buyel, 2019). In this context, when confronted with the CoVid-19 crisis, we decided to exercise a partial reorientation of the activities in our academic lab, which is equipped with a multipurpose glass greenhouse facility, towards the production of SARS-CoV-2 antigens and antibodies against the virus. Here we describe the recombinant production, purification and analysis of six anti-SARS-CoV-2 monoclonal antibodies at laboratory scale, plus a pilot upscaling of two of those six antibodies. Next to production scale, a critical parameter to assess was the response time. The process described here started in mid-April 2020 with the selection of literature-available antibody variable sequences and finalized nine weeks later with approximately 0.2 g of anti-SARS-CoV-2 antibody (Ab) produced in modular badges of 56 *N. benthamiana* plants and formulated as one litre of Ab-enriched plant apoplastic fluid. Based on this experience, we estimate that the same process can be reduced up to 6-7 weeks with small pre-adaptations, a remarkably short reaction time for a *de novo* antibody production system. Absolutely key for this fast reaction is the immediate availability of scientific data including antibody sequences in pre-print repositories. This is in our opinion one of the most positive lessons that can be extracted from the CoVid-19 crisis. We discuss here the possible applications of the fast plant-produced antigens and antibodies in diagnostics and therapy and propose the repurpose of high-tech agricultural facilities as an alternative for decentralized biomanufacturing in times of crisis.

## MATERIALS AND METHODS

### Plant material and growth conditions

Both, *Nicotiana benthamiana* wild type plants and 1,2-xylosyltransferase/alpha1,3-fucosyltransferase (ΔXT/FT) RNAi knock down lines (Strasser et al., 2008) were grown in the greenhouse. Growing conditions were 24 °C (light)/20 °C (darkness) with a 16-h-light/8-h-dark photoperiod.

### Cloning of anti-SARS-CoV-2 antibodies and SARS-CoV-2 antigens

All sequences were cloned and assembled using the GoldenBraid (GB) assembly system (Sarrion-Perdigones et al., 2013; https://gbcloning.upv.es). Antibody sequences were obtained from literature (see Table 1). All antibodies were cloned as single chain antibodies fused to the human IgG1 Fc domain. Those antibodies derived from synthetic or camelid single domain VHH libraries (sybody 3, sybody 17 and nanobody 72) were designed as direct fusions. CR3009, CR3018 and CR3022 human monoclonal antibodies were redesigned as single chain variable fragment (ScFv) by connecting the variable light (VL) and heavy (VH) chains with a GGGGSGGGGSGGGGSSGGGS peptide linker. Antibody sequences were codon optimised for *N benthamiana* with the IDT optimization tool at http://eu.idtdna.com/CodonOpt.

**Table 1.**
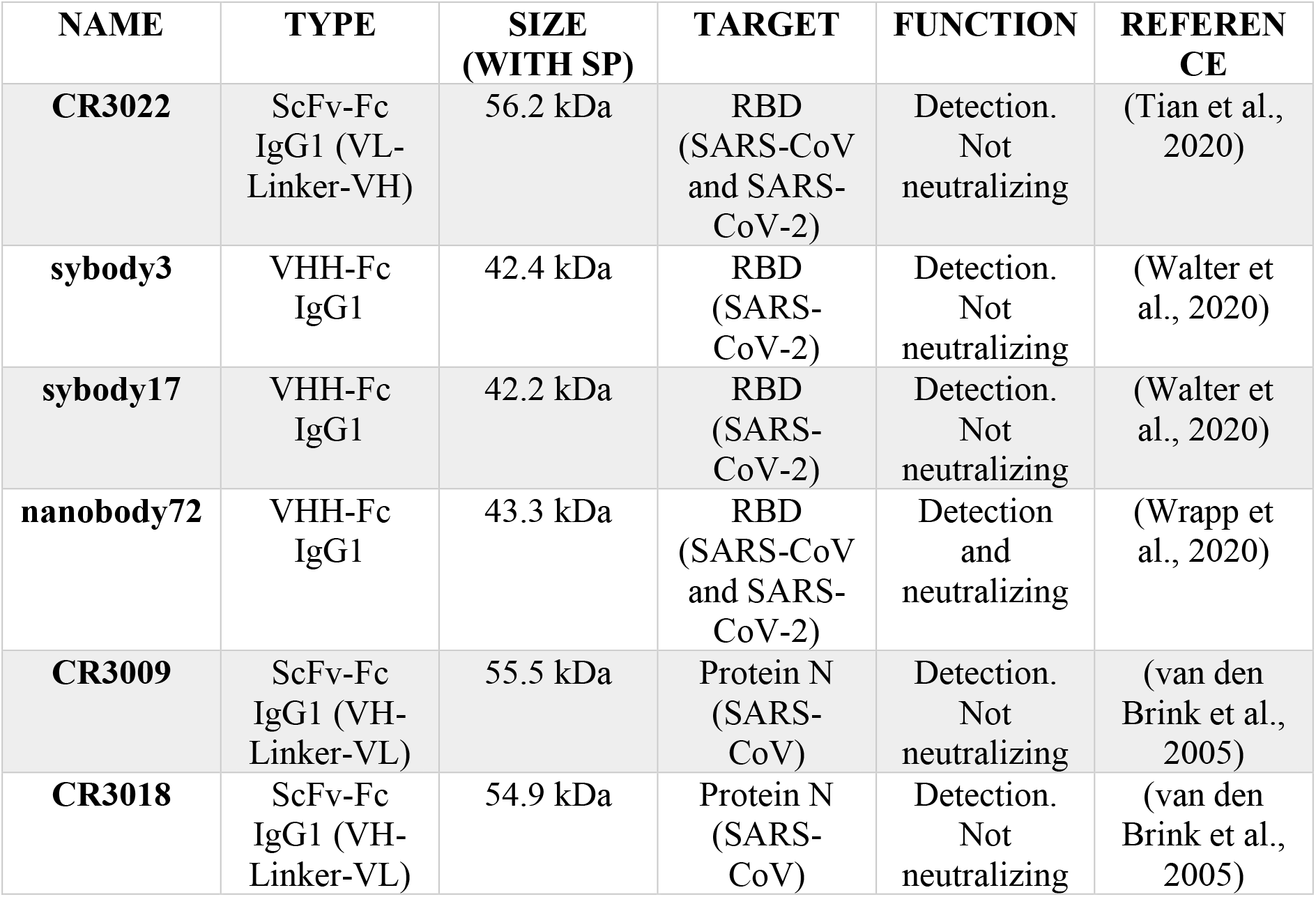
Detailed information of six different antibodies selected for recombinant expression in plants.

| NAME | TYPE | SIZE<br>(WITH SP) | TARGET | FUNCTION | REFEREN<br>CE |
| --- | --- | --- | --- | --- | --- |
| **CR3022** | ScFv-Fc<br>IgG1 (VL-Linker-VH) | 56.2 kDa | RBD<br>(SARS-CoV<br>and SARS-CoV-2) | Detection.<br>Not<br>neutralizing | (Tian et al.,<br>2020) |
| **sybody3** | VHH-Fc<br>IgG1 | 42.4 kDa | RBD<br>(SARS-CoV-2) | Detection.<br>Not<br>neutralizing | (Walter et<br>al., 2020) |
| **sybody17** | VHH-Fc<br>IgG1 | 42.2 kDa | RBD<br>(SARS-CoV-2) | Detection.<br>Not<br>neutralizing | (Walter et<br>al., 2020) |
| **nanobody72** | VHH-Fc<br>IgG1 | 43.3 kDa | RBD<br>(SARS-CoV<br>and SARS-CoV-2) | Detection<br>and<br>neutralizing | (Wrapp et<br>al., 2020) |
| **CR3009** | ScFv-Fc<br>IgG1 (VH-Linker-VL) | 55.5 kDa | Protein N<br>(SARS-CoV) | Detection.<br>Not<br>neutralizing | (van den<br>Brink et al.,<br>2005) |
| **CR3018** | ScFv-Fc<br>IgG1 (VH-Linker-VL) | 54.9 kDa | Protein N<br>(SARS-CoV) | Detection.<br>Not<br>neutralizing | (van den<br>Brink et al.,<br>2005) |

The SARS-CoV-2 antigen sequences used (N protein, YP_009724397.2; and S-protein RBD domain, YP_009724390.1, aa 319-541) derive from the Wuhan strain NC_045512. Four different versions of RBD were designed corresponding to (i) the native sequence with a C-terminal 6xHis-Tag or (ii) an N-terminal 6xHis-Tag and a C-terminal KDEL sequence for ER retention, and (iii and iv) their corresponding *N. benthamiana* codon optimised counterparts, using the same tool as above.

DNA sequences were domesticated as level 0 phytobricks for GB cloning and ordered for synthesis as double-stranded DNA fragments (gBlocks, Integrated DNA Technologies). gBlocks were first cloned into the domestication vector pUPD2 (Vazquez-Vilar et al., 2017) in a BsmBI Golden Gate restriction/ligation reaction (37 °C - 10 min, 50x (37 °C - 3 min / 16 °C - 4 min), 50 °C - 10 min, 80 °C - 10 min). The ligation product was transformed into *E. coli* Top 10 electrocompetent cells and positive clones were verified by restriction digestion analysis and sequencing. pUPD2 level 0 phytobricks were then cloned into the expression vectors pGreen SP-hIgG1 (antibody sequences), pCambiaV1 (RBD sequences) or pCambiaV2 (N sequences). pGreen SP-hIgG1 is a pGreen vector based adaptation of the the MagnICON® 3’ provector pICH7410 (ICON Genetics) that is designed for BsaI cloning of GB (B4-B5) standard parts as in-frame fusions with the tobacco (1-3)-beta-glucanase signal peptide and the human IgG1 Fc domain. Similarly, pCambiaV1 and pCambiaV2 are pCambia based adaptations of the MagnICON® 3’ provector pICH7410 that are designed for BsaI cloning of GB standard parts as in-frame fusions with tobacco (1-3)-beta-glucanase signal peptide (pCambiaV1, for expression of secreted proteins) or without any subcellular localization signal (pCambiaV2, for expression of cytoplasmic proteins). Assembly reactions were performed as above, and the ligation reactions were transformed into *E. coli* Top 10 electrocompetent cells. Positive clones were verified by restriction digestion analysis. All level 0 parts generated in this work are listed in Supplementary Table 1.

### Transient expression in *Nicotiana benthamiana*

For transient expression in *N. benthamiana,* the plasmids were transformed into *Agrobacterium tumefaciens* strain *GV3101 C58C1* by electroporation. The same strain but carrying the pSoup helper plasmid was employed to allow the replication of the pGreen vectors which encode the antibodies. Overnight grown exponential cultures were collected by centrifugation and the bacterial pellets were resuspended in agroinfiltration solution (10 mM MES, 20 mM MgCl2, 200 μM acetosyringone, pH 5.6) and incubated for 2 h at RT in a horizontal rolling mixer. For small scale agroinfiltration, culture optical density at 600 nm was adjusted to 0.1 with agroinfiltration solution and the bacterial suspensions harbouring the 3’ antibody or antigen modules, the Integrase (pICH14011), and the 5’ module (pICH17388) were mixed in equal volumes. Control samples were agroinfiltrated with pICH11599_DsRed and Integrase module. Agroinfiltration of 5 to 6-week-old *N. benthamiana* plants was carried out through the abaxial leaf surface using a 1 ml needle-free syringe (Becton Dickinson S.A.). For pilot scale production, the bacterial suspensions were prepared as above except that a lower OD600 was used (0.005 for sybody17 agroinfiltration and 0.01 for nanobody72 agroinfiltration). Additionally, for sybody17 agroinfiltration, a bacterial suspension of pICH11599_DsRed, a MagnICON® 3’module encoding the fluorescent protein DsRed, was added to the final *Agrobacterium* infiltration solution in a ratio 1:1:0.9:0.1 (pICH17388:pICH14011:pGreenSP-Sybody17-hIgG1:pICH11599_DsRed). Delivery of *Agrobacterium* to the plant cells was carried out by vacuum infiltration in a vacuum degassing chamber (model DP118, Applied Vacuum Engineering) provided with a 30 L infiltration tank. The aerial part of whole plants (seven plants at a time) was immersed into the *Agrobacterium* infiltration solution; vacuum was applied for 1 min at a vacuum pressure of 0.8 bar and then slowly released.

### Apoplast fluid extraction

14 days post-vacuum agroinfiltration leaves were excised and then infiltrated with 20 mM Phosphate buffer (7.4 mM NaH2PO4, 12.6 mM Na2HPO4.7H2O, pH 7), without (sybody17) or with (nanobody72) 0.5 mM PMSF (Sigma-Aldrich, #78830), following the same procedure as the vacuum agroinfiltration. After eliminating the buffer excess with tissue paper, the leaves were introduced into mesh zipped bags and then centrifuged using a portable cloth dryer *Orbegozo SC4500.* Thus, the apoplastic fluid was obtained from the drain tube. The apoplastic fluid was centrifuged (10 min, 11000 x *g*, at 4 °C) to remove any cell debris and *Agrobacterium,* the supernatant was collected and then 4 fractions of 4mL were concentrated 8 times using 10 kDa *Amicon Ultra-4 10K Centrifugal filters* (Millipore) after centrifugation (20 min, 3700 x *g,* at 4 °C).

### Antibody extraction and purification

Protein crude extracts were obtained by homogenizing ground frozen leaf tissue with cold PBS buffer (20 mM NaH2PO4, 80 mM Na2HPO4.7H2O, 100 mM NaCl, pH 7.4) in a 1:3 (w/v) and were centrifuged at 13000 rpm for 15 min at 4 °C. For antibody purification, 4 g of ground agroinfiltrated tissue were extracted in 12 mL of cold 20 mM Phosphate buffer. Samples were centrifuged at 10000 x *g* for 15 min and the supernatant was transferred to a clean tube and further clarified by filtration through a 0.22 μm membrane filter. The recombinant antibodies were purified by affinity chromatography with Protein A agarose resin (ABT Technology) following a gravity-flow procedure according to the manufacturer’s instructions. 100 mM citrate buffer pH 3 was used for elution and 1 M Tris-HCl pH 9 was used for neutralization of the eluted sample (37.5 μL for each 250 μL elution fraction). Purified antibodies were quantified using the Bio-Rad Protein Assay following the manufacturer’s instructions and using BSA for standard curve preparation.

### Antigen extraction and purification

The *N. benthamiana* leaves infiltrated with the different SARS-CoV-2 proteins were collected 5 (RBD) or −7 (N protein) days post infiltration (dpi). Leaves were frozen in liquid nitrogen and stored at −80 °C until used.

Protein extraction was performed using 3-6 g of ground frozen tissue in 3 volumes of cold-extraction buffer. Three different buffers were tested as a first approach, in order to optimize the purification yields. Buffer A: PBS buffer with 10 mM imidazole, pH 8. Buffer B: Buffer A supplemented with 1% Triton X-100, and Buffer C: Buffer B supplemented with 20% glycerol, 10% sucrose and 0.05% 2-β-mercaptoethanol. Samples were vigorously vortexed and centrifuged at 10000 x *g* for 15 min at 4 °C. The supernatant was carefully transferred to a clean tube and filtered through a 0.22 μm syringe filter. Protein purification was carried out by Ni-NTA affinity chromatography as described in (Fernandez-del-Carmen et al., 2013). Purified proteins were quantified using the Bio-Rad Protein Assay following the manufacturer’s instructions and using BSA for standard curve preparation.

### SDS-PAGE and Western Blot Analysis

Proteins were separated by SDS-PAGE electrophoresis on NuPAGE 10% Bis-Tris Gels (Invitrogen) using MES-SDS running buffer (50 mM MES, 50 mM Tris-base, 3.5 mM SDS, 1 mM EDTA, pH 7.3) under reducing conditions. Gels were visualized by Coomassie blue staining. For Western blot analysis, proteins were transferred to PVDF membranes (Amersham Hybond™-P, GE Healthcare) by semi-wet blotting (XCell II™ Blot Module, Invitrogen, Life Technologies) following the manufacturer’s instructions. Blots were blocked with 2% ECL Prime blocking agent (GE Healthcare) in PBS-T (PBS buffer supplemented with 0.1% (v/v) Tween-20). For anti-SARS-CoV-2 antibody detection, the blots were incubated with 1:20000 HRP-conjugated rabbit anti-human IgG (Sigma-Aldrich, #A8792). For SARS-CoV-2 antigen detection the blots were incubated with 1:2000 Anti-His mouse monoclonal primary antibody (Qiagen, #34660) and then incubated with 1:10000 peroxidase labelled anti-mouse IgG secondary antibody (GE Healthcare). Blots were developed with ECL Prime Western blotting Detection Reagent (GE Healthcare) and visualised using a Fujifilm LAS-3000 imager.

### Antigen ELISA

The overnight coating of Costar 96 Well EIA/RIA plates (Corning) was carried out at 4 °C with 100 μL of 4 μg/mL RBD (RayBiotech, #230-30162) or BSA (used as control) in Coating buffer (15 mM Na2CO3, 35 mM NaHCO3, pH 9.8). After 4 washes with 300 μL of PBS, the plate was blocked with 200 μl of a 2% (w/v) ECL Advance Blocking Reagent (GE Healthcare) solution in PBS-T (PBS supplemented with 0.1% (v/v) Tween-20) for 2 h at RT. The plate was washed 4 times with PBS, and then, starting at 2 μg of the purified antibody per well (100 μl), 1:5 serial dilutions in blocking solution were incubated for 1 h 30 min at RT. After 4 washing steps with PBS-T (PBS buffer supplemented with 0.05% Tween-20), 1:2000 HRP-labelled rabbit anti-human IgG (Sigma-Aldrich, #A8792) in blocking solution was added. After 1 h, the plate was washed with PBS and the substrate o-phenylenediamine dihydrochloride SIGMAFAST™ OPD tablet (Sigma-Aldrich, #P9187) was added (following manufacturer’s instructions). Reactions were stopped with 50 μL 3 M HCl per well and absorbance was measured at 492 nm. The endpoint titer was determined as the last concentration of each purified antibody showing an absorbance value higher than the value defined as cutoff (mean blank + 3SD). Blank is defined as the values from each ELISA test against BSA (Zrein et al., 1986; Armbruster and Pry, 2008).

### Sandwich ELISA

The sandwich ELISAs were performed as described in the antigen ELISA section with a few changes. The plates were coated with 100 μL of 4 μg/mL murine anti-His mAb (Qiagen, #34660), and after blocking, the plates were incubated with 100 μl of the crude extracts of the (RBD/N) antigen expressing leaves serially diluted (1:2) in BSA 0.5%. WT crude extracts were used as negative control. The crude extracts were prepared by adding a volume of PBS buffer corresponding to 3 times the mass of the ground tissue in liquid nitrogen. Then the mix was centrifuged (13000 rpm, 4 °C, 15 min) and the supernatant was subjected to sonication before use. The antigens were sandwiched with 1 μg of the corresponding purified antibody (or 100 μL of the apoplastic fluid in 2% blocking reagent) per well (1 h 30 min incubation, RT). The same procedure as in the antigen ELISA was followed for the incubation with the conjugated secondary antibody, colorimetric reaction and measurement.

## RESULTS

### Cloning and expression of anti-SARS-CoV-2 recombinant antibodies

Six different antibody sequences were selected for recombinant production in *N. benthamiana,* following a plant deconstructed viral strategy based on Magnifection technology, as described earlier (Marillonnet et al., 2005) (see Table 1). Four of those were directed against the Receptor Binding Domain (RBD) of the SARS-CoV-2 spike (S) protein, whereas the remaining two were directed against the N protein. All six antibodies were engineered as single polypeptide chains fused to the human Cγ2-Cγ3 constant immunoglobulin domains. Three of them, those derived from single chain camelid or synthetic VHH antibody libraries, were produced as direct fusions. The other three, derived from full-size human monoclonal antibodies, were redesigned as scFvs, using a linker peptide that connects VH and VL regions (see Fig 1A). The nucleotide sequences of the different variable regions were all obtained from the literature, then chemically synthesized with appropriate extensions and cloned into a destination Magnifection-adapted vector using a type IIS restriction enzyme strategy. The cloning cassette was flanked by a β-endoglucanase signal peptide for apoplastic localization in N-terminal and the human Cγ2-Cγ3 domains of the human IgG1 in the C-terminal side. The resulting vectors were transferred to *Agrobacterium* cultures and agroinfiltrated in *N. benthamiana* leaves in combination with a 5’ MagnICON® module, containing the RNA polymerase and movement protein, and with an integrase module (Fig 1B). For antibody production we used wild-type and RNAi ΔXT/FT glycoengineered *N. benthamiana* plant lines, the latter lacking plant-specific Xylose and Fucose glycosylation (Strasser et al., 2008). Infiltrated leaves were examined daily, and only minimal damage was observed in the agroinfiltrated tissues during the incubation period. After seven days, leaf samples were collected, ground, and crude extracts were analyzed in SDS-PAGE. As can be observed in Fig 2A-B (upper panel), all samples produced Coomassie-detectable bands of the expected antibody size. scFv-IgG1 55-56 kDa antibodies migrated slightly above the 50-55 kDa Rubisco large subunit, partially masking its detection. VHH-IgG1 antibodies migrated at the expected 42-43 kDa size. The identity of the Coomassie bands was confirmed by Western blot using an anti-human IgG1 antibody for detection (Fig 2A-B, lower panel). As shown in Fig 2A-B, under reducing conditions lower molecular weight (MW) bands were also detected, probably as a result of partial proteolytic degradation. Small-scale affinity purification was carried out for all six antibodies produced in ΔXT/FT plants using protein A affinity chromatography (Fig 2C-D). The resulting purified antibodies were used to estimate the yield of the final product, which ranged between 73.06 μg/g FW (CR3022 antibody) to 192.63 μg/g FW (nanobody72 antibody) (see Table 2).

**Figure 1.**
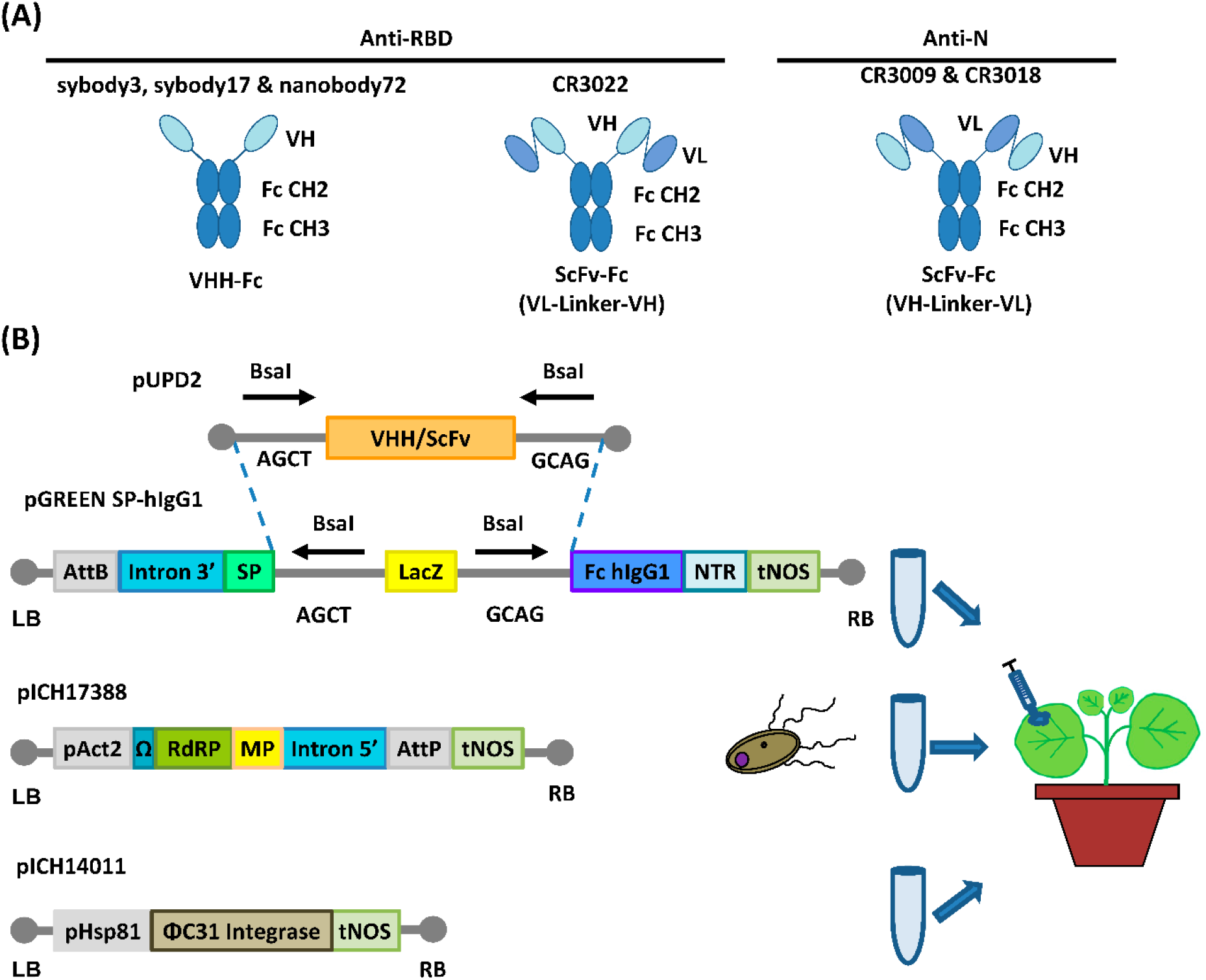
Design for the expression of anti-SARS-CoV-2 recombinant antibodies. **(A)** Schematic representations of VHH-Fc and ScFv-Fc antibody versions. VHH, heavy chain only single domain antibody; ScFv, single chain variable fragment; Fc, human IgG1 Cγ2-Cγ3 domains. **(B)** Schematic representation of the cloning procedure for transient expression via MagnICON®. VHHs and ScFvs cloning into the pGreen SP-hIgG1 vector and coinfiltration together with the plasmids pICH17388 and pICH14011 for the *in planta* viral reconstruction (through ΦC31 mediated PB recombination) and antibody expression. AttB & AttP, site-specific recombination site; SP, signal peptide; NTR, nontranslated region; tNOS, terminator of nopaline synthase; pAct2, actine2 promoter of *Arabidopsis thaliana;* RdRP, RNA-dependent RNA polymerase; MP, viral movement protein; pHsp81 promoter of the heat shock protein 81.

**Figure 2.**
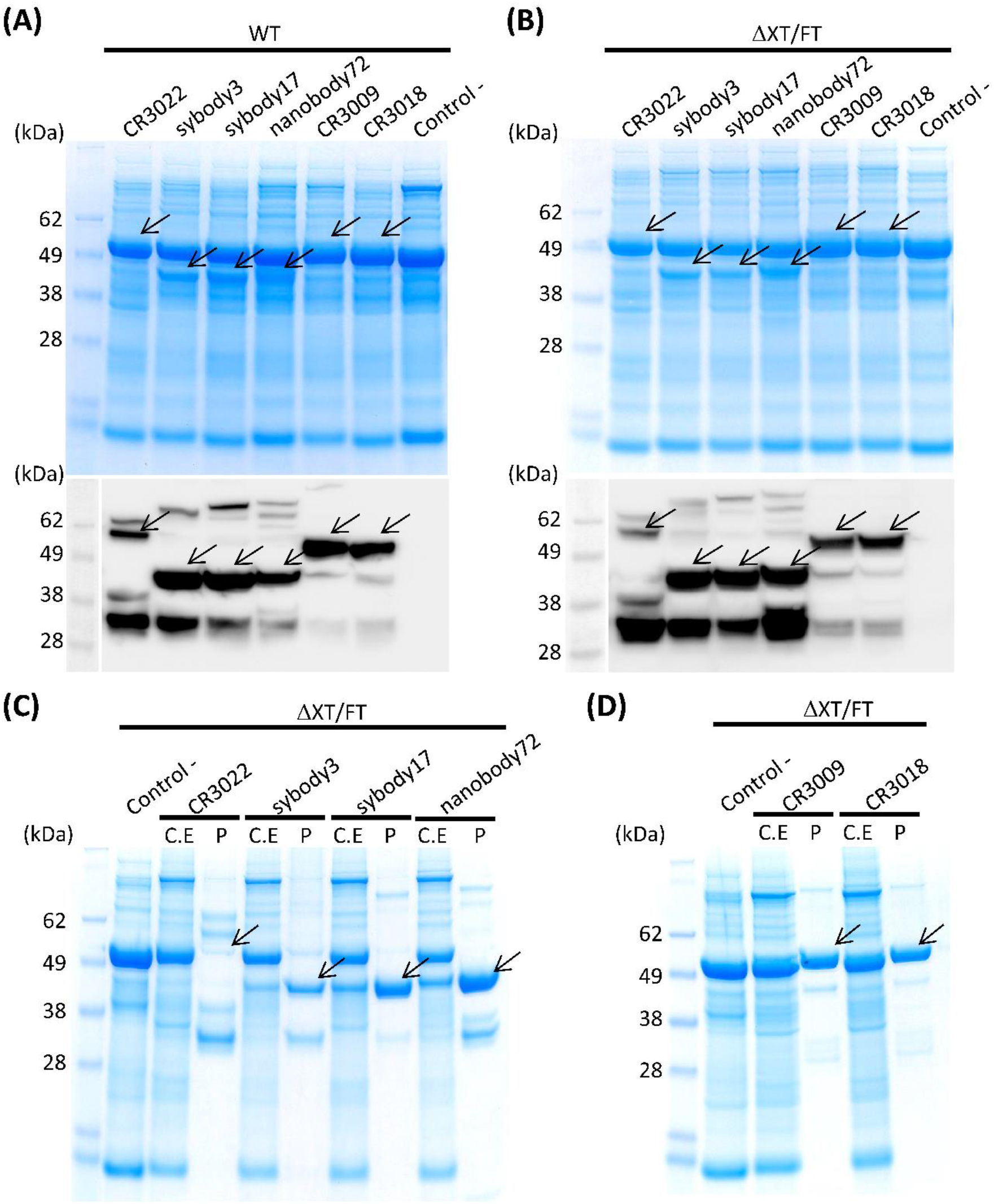
Analysis of the expression of all six anti-SARS-CoV-2 recombinant antibodies produced in *N. benthamiana* leaves. Coomassie stained SDS gels (upper panel) and Western blots (down panel) of protein extracts from wild-type **(A)** and ΔXT/FT **(B)** *N.benthamiana* leaves agrotransformed all of them with six antibodies. 17 μl of crude extract were loaded per well in SDS gels. The blots were incubated with an antibody against constant region of human IgG. **(C),(D)** Coomassie stained SDS gels with purified (P) recombinant antibodies from ΔXT/FT plants compared to their corresponding protein crude extract (C.E) for each analyzed antibody. 20 μg of total protein from crude extract per well and 3 μg of purified protein per well were loaded in SDS gels. Control samples (-) were agroinfiltrated as indicated in Methods. Arrows indicate the presence and position of the corresponding band protein for each antibody.

**Table 2.** Yield estimation for all six purified recombinant antibodies, RBD and N proteins.

| Purified<br>protein | Yield<br>(µg/g FW) | % TSP |
| --- | --- | --- |
| **CR3022** | 73.06 | 1.96 |
| **sybody3** | 122.53 | 2.56 |
| **sybody17** | 153.36 | 3.31 |
| **nanobody72** | 192.63 | 3.18 |
| **CR3009** | 73.38 | 1.88 |
| **CR3018** | 81.24 | 1.79 |
| **nRBD:His (Buffer A)** | 4.31 | 0.18 |
| **nRBD:His (Buffer B)** | 4.02 | 0.051 |
| **bRBD:His (Buffer A)** | 2.94 | 0.097 |
| **bRBD:His (Buffer B)** | 5.21 | 0.051 |
| **His:bN (Buffer A)** | 30.98 | 0.45 |

### Cloning and expression of SARS-CoV-2 recombinant antigens

The *in-planta* production of SARS-CoV-2 RBD and N protein antigens was also assayed in parallel using a similar strategy as described for antibody production. For this purpose, two versions of the expression vector were designed for RBD, one with the native viral sequence and the other with a *N. benthamiana* codon-optimized sequence. For the N protein, only the *N. benthamiana* codon-optimized sequence was employed. For RBD, native and codon optimized versions were targeted to the apoplast with the tobacco glucan endo-1,3-beta-glucosidase signal peptide and versions containing a KDEL peptide for ER retention were also generated. All nucleotide sequences were chemically synthesized with a small nucleotide extension coding for a Histidine tag for detection (Fig 3A).

**Figure 3.**
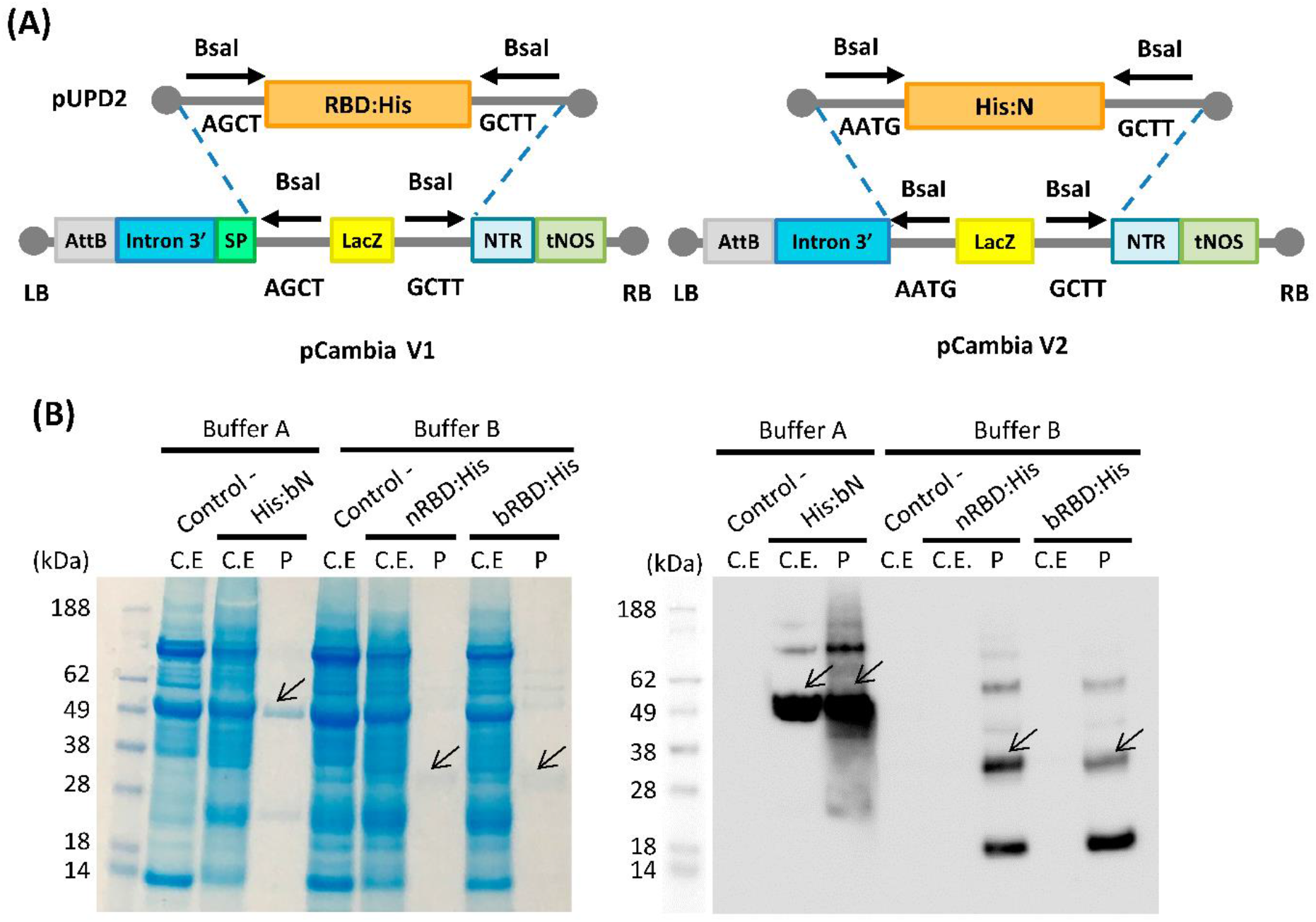
SARS-CoV-2 RBD and N proteins expression in *N. benthamiana* agroinfiltrated leaves. **(A)** Schematic representation of the pUPD2 and 3’ module vectors generated for antigen production. **(B)** Coomassie stained SDS-PAGE (left) and Western blot (right) of crude extracts from non-infiltrated *N. benthamiana* leaves (Control -), leaves infiltrated with viral vectors for the expression of His:bN, nRBD:His and bRBD:His and the corresponding purified proteins.

As described for antibody production, MagnICON®-derived 3’ vector modules encoding RBD and N proteins were agroinfiltrated in combination with an integrase module and a 5’-module lacking any additional subcellular localization signal. Shorter incubation times were decided in antigen production as compared to antibodies because antigen constructs produced different degrees of necrotic lesions in the leaves, ranging from mild symptoms in N protein to severe necrosis after four days in native RBD. For those constructs producing more severe lesions, incubation time was reduced to five days, and for the rest the incubation period was extended to seven days. RBD was extracted and purified using small-scale affinity-chromatography with Niquel columns and the resulting Coomassie and a Western blot analysis are shown in Fig 3B. RBD can be detected as a major estimated 30 kDa band, with the presence of higher MW bands that suggest multimerization. ER retention did not improve expression levels of RBD for the native version, nor for the *N. benthamiana* optimized one (data not shown). Addition of 1% Triton X-100 to the standard extraction buffer (see Materials and Methods) did not improve the yield, which was estimated as 2-4 μg/g FW (Table 2). N protein was extracted from agroinfiltrated leaves and affinity purified following the same procedure described for RBD. A major 49 kDa band was detected both on the crude extract and upon purification (Fig 3B). Small-scale affinity-chromatography with Niquel columns gave an estimated yield of 30 μg/g FW for N protein (Table 2).

### Characterization of antigen-antibody binding activities

Binding activities of affinity purified anti-RBD antibodies were analysed by antigen ELISA as shown in Fig 4A. As expected, all assayed antibodies were active in binding their respective antigen. Endpoint dilution titers were calculated for anti-RBD antibodies using a commercial antigen. Sybodies 3 and 17 and nanobody72 showed high dilution titers, (0.75 μM, 0.03 μM and 0.03 μM, respectively), but the performance of CR3022 was significantly lower (2.85 μM). In a parallel experiment, we tested the ability of plant-made antibodies to selectively detect our own plant-made antigens, including here also the N protein, using a sandwich ELISA approach. For this analysis, ELISA plates were coated with a murine anti-His mAb, incubated with serial dilutions of crude plant extracts from antigen-producing plants and sandwiched with purified plant-made antibodies. As shown in Fig 4B-C, all antigen-producing plant extracts gave sandwich-ELISA signals significantly above the background when assayed using their cognate antibodies, thus evidencing the capacity of both, antibodies and antigens, to function as potent diagnostic tools. Background signals in this experiment are likely due to cross-reaction of the anti-human IgG secondary antibody with the murine anti-His mAb, and could be easily reduced for more potent diagnostic applications by employing recombinant antibody formats other than IgG.

**Figure 4.**
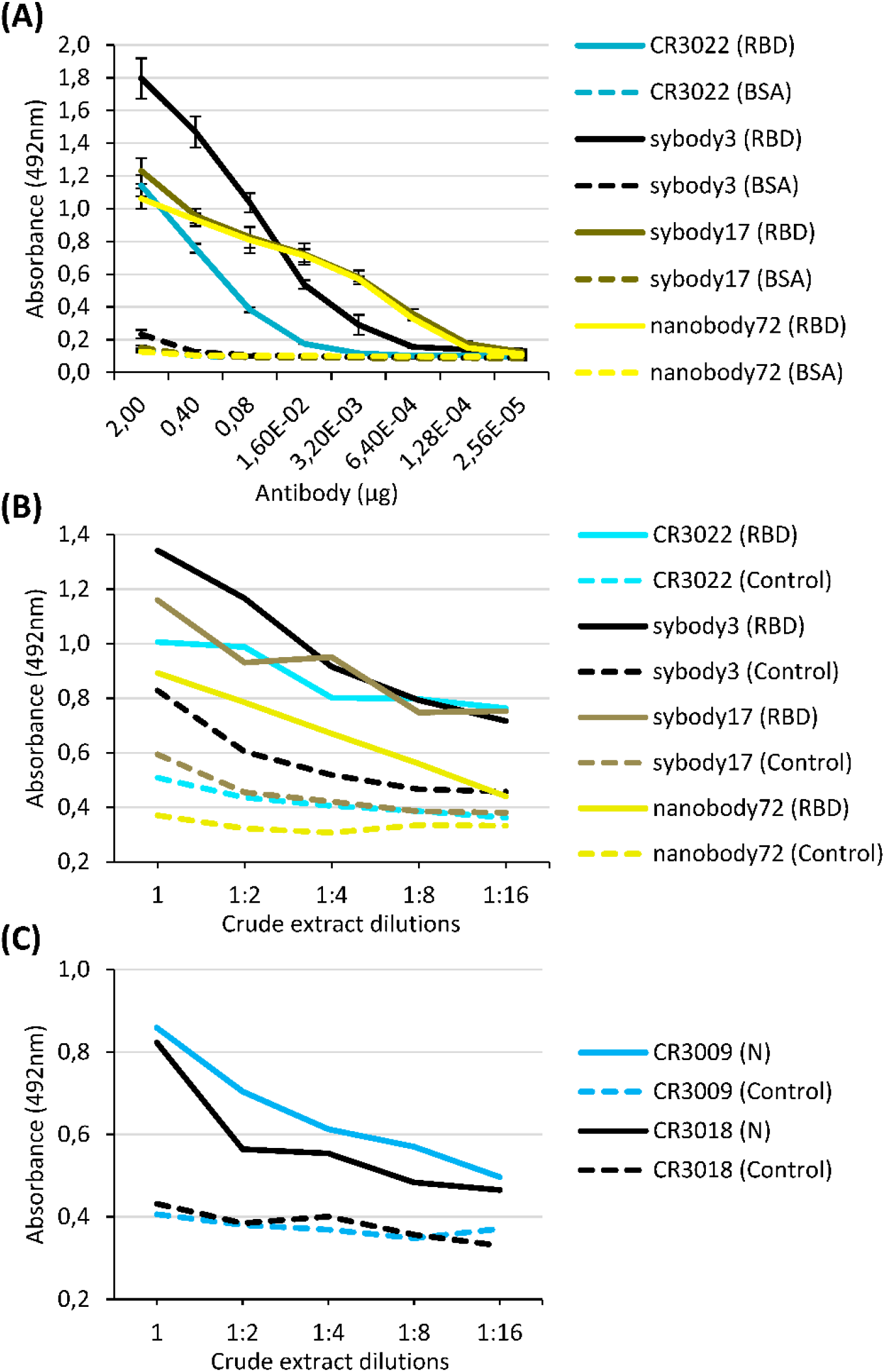
Comparison of binding activities of anti-SARS-CoV-2 purified antibodies by ELISA tests. **(A)** Antigen ELISA titer graph of serial dilutions 1:5 of anti-RBD purified antibodies from an initial amount of 2 μg per well. The titer was calculated by absorbance measurements at 492 nm. The X axis indicates the amount (μg) of antibody per well. Solid lines indicate the ELISA test against RBD and dashed lines indicate the corresponding control ELISA tests against BSA. Bars represent mean ± SD, n = 3 independent replicates. **(B),(C)** Sandwich ELISA titer graph with anti-RBD **(B)** and anti-N **(C)** purified antibodies (1 μg per well). Serial dilutions 1:2 of protein crude extracts of antigens (RBD/N) expressed in plants were used to test the affinity of each antibody. Wild type crude extracts samples were used as negative control. Solid lines indicate the ELISA test against RBD and N antigens and dashed lines indicate the corresponding control ELISA tests.

### Pilot antibody upscaling and analysis of apoplastic fluid

In the design of a pilot upscaling experiment, we favoured modularity and tried to maximize the affordability and adaptability of the process by reducing the requirements for highly specialized lab equipment. We carried out a final agroinfiltration for recombinant antibody production using a total of 112 plants, equivalent to approximately 2.5 kilograms of fresh plant material. The plants were divided in two batches of 56 plants each, and used to produce sybody17 and nanobody72 respectively, as these antibodies showed the most promising binding activities and yields. To facilitate the upscaling of the agroinfiltration process, plant seedlings were transplanted in growth modules, each module comprising seven pots kept together in a double layer of disposable plastic-board hexagons as shown in Fig 5A. Each production batch consisted in eight hexagonal modules. When plants were six weeks old, they were agroinfiltrated by submerging each hexagon upside down into a 40 cm diameter cylindrical tank filled with 30 L of an *Agrobacterium* suspension, set inside a cylindrical vacuum degas chamber (Fig 5A). In this way, seven plants at a time were vacuum-agroinfiltrated by slowly releasing vacuum while leaves remained submerged in the solution. Next, plants were rinsed, brought back to the growth chamber and incubated for 14 days before harvest. Two different concentrations of the *Agrobacterium* suspension were used in this experiment. One of them (sybody17) consisted in an OD600 0.005-final mix containing plasmids pICH14011, pICH17388, pGreenSybody17-IgG1, and pICH11599_DsRed at 1:1:0.9:0.1 ratio, where pICH11599_DsRed is a MagnICON® 3’module encoding DsRed. The fluorescent marker was added to the infiltration mix to monitor the extension of the viral infection foci. As described elsewhere (Julve et al., 2013; Julve Parreño et al., 2018) superinfection exclusion among virial clones yields mosaic-like expression patterns of individual clones, therefore the tiles produced by red fluorescent proteins were used as an indication of the extension and distribution of the unlabelled foci producing the recombinant antibody. In parallel, nanobody72 upscaled production was undertaken by agroinfiltration of an OD600 0.01 *Agrobacterium* culture mix containing pICH14011, pICH17388 and pGreenNanobody72-IgG1 at 1:1:1 ratio. After 14 days, DsRed tiles in sybody17 experiment, clearly visible with the naked eye, finalized their expansion in most agroinfiltrated leaves, an indicator that the expression tiles had covered the whole leaf surface (Fig 5B). At this stage, leaves were harvested and submitted to an apoplastic fluid recovery assay, where >0.5 Kg batches of detached leaves were vacuum infiltrated in 20 mM Phosphate buffer using the same vacuum device as described above. Once rinsed to remove the excess of buffer, leaves were packed in mesh zipped bags, spinned down in a spin portable cloth dryer, and the intercellular apoplastic fluid was recovered from the drain tube. With this simple procedure, between 940 and 1200 millilitres of apoplastic fluid (sybody 17 and nanobody 72, respectively) was recovered from 1.2 Kg of detached leaves. A fraction of the apoplastic fluid of both antibodies was concentrated 8 times in 10 kDa centricons, and the rest was kept refrigerated for further analysis. Fig 5C-D show the Coomassie-staining and western blot analysis of crude extracts as well as apoplastic fluid preparations, and their corresponding purifications. Crude extracts in this pilot experiment showed a VHH-IgG1 band similar in intensity to that obtained in small scale experiments (data not shown). Interestingly, apoplastic fluid consisted in a very simplified mix of proteins, with the recombinant antibody being among the most predominant ones. As shown, the different optical density of the *Agrobacterium* culture, together with the presence of a competing DsRed clone clearly influenced the accumulation levels, with the yields of nanobody72 clearly outperforming those of its sybody counterpart. Unfortunately, the antibodies seemed partially degraded as indicated by the presence of two bands smaller than the expected VHH-Cγ2-Cγ3 size, which could be compatible with degradation fragments. Degradation was only partially solved with the addition of the protease inhibitor PMSF into the recovered Phosphate buffer, as shown with nanobody72 production (Fig 5D). Despite degradation, in a densitometric analysis we estimate that the recovered apoplastic fluid contains 0.2 g per liter of intact mAb full-size. Finally, we performed sandwich ELISA tests of sybody17 and nanobody72 (Fig 5E and Fig 5F, respectively) using the total and concentrated apoplastic fluid as detection reagent against serial dilutions of crude plant extracts from RBD-producing plants, showing that this simple antibody preparation can be directly employed in detection procedures without the need of additional purification steps.

**Figure 5.**
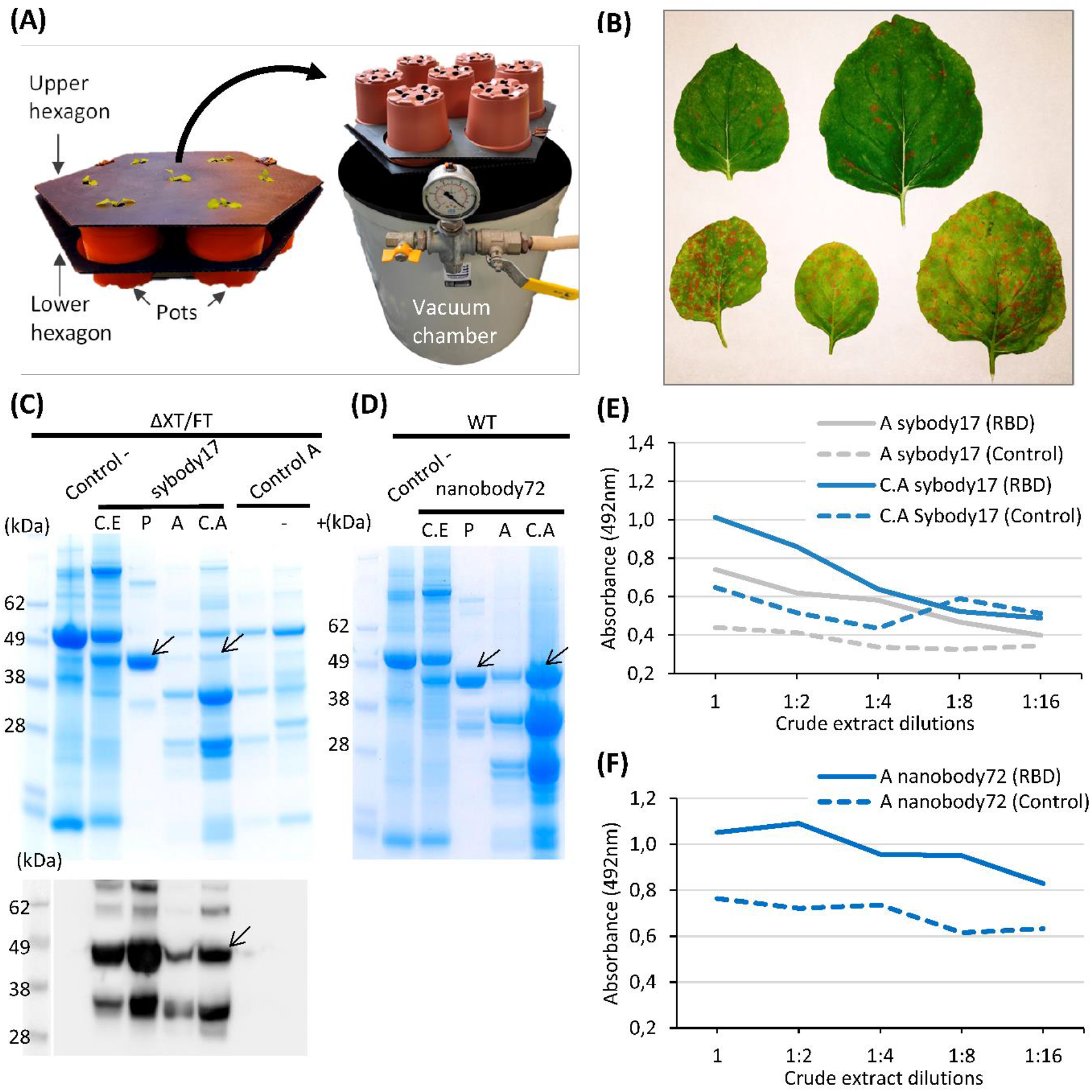
Analysis of upscaling production of anti-RBD sybodyl7 and nanobody72 antibodies as an example for a pilot assay. **(A)** Hexagon growth modules used in the pilot assay with *N. benthamiana* plants and vacuum degas chamber used for vacuum agroinfiltration. **(B)** Different agroinfiltrated ΔXT/FT *N. benthamiana* leaves with DsRed tiles as indicator of the expression of the recombinant antibody. **(C),(D)** Coomassie stained SDS gel and Western blot of protein extracts from ΔXT/FT plants expressing sybody17 antibody **(C)** and Coomassie stained SDS gel of protein extracts from WT plants expressing nanobody72 antibody **(D);** from total crude extract (C.E), purified extract (P), total apoplastic fluid (A) and centricon-concentrated apoplastic fluid (C.A). Wild type crude extracts (control -) and apoplastic fluid with (+) or without (-) DsRed expression were used as control samples. 17 μl of total protein extract per well and 3 μg of purified protein per well were loaded in SDS gels. Arrows indicate the presence and position of the corresponding band protein for each antibody. **(E),(F)** Sandwich ELISA tests with total and/or concentrated apoplastic fluid extract from plants expressing sybody17 **(E)** or nanobody72 **(F)**. Serial dilutions 1:2 of protein crude extracts of RBD expressed in plants was used to test the affinity (indicated by solid lines). Wild type crude extracts samples were used as negative control (indicated by dashed lines).

## DISCUSSION

Several *N. benthamiana*-dedicated bioproduction facilities are functioning worldwide, as those from Leaf Expression Systems in UK (Dobon, 2019), Icon Genetics (Giritch et al., 2006) and Fraunhofer in Germany (Wirz et al., 2012) or Kentucky Bioprocessing in US (Olinger et al., 2012), among others. Notably, Medicago recently announced the building a new 44000 sqm facility with capacity for around 40-50 million of planned doses of flu vaccine per year. Such facilities usually involve separated modules for upstream processing, namely a wet-lab module for preparation of the bacterial inoculum, a regular plant growth chamber, and agroinfiltration room, and a post-infiltration growth chamber. In addition, downstream processing facilities are often situated next or to the production ones to minimize the handling time of fresh tissues. Whereas installed capacity of plant-dedicated biofactories is in continue growth, they are clearly insufficient to respond to global or even regional demands in times of crisis. We reasoned that, at least for upstream processes, the infrastructures required for medium scale *N. benthamiana*-based production are not radically different to those employed in high-tech agriculture practices as hydroponics, aeroponics or vertical farming, and thus high-tech agriculture facilities could be easily repurposed as biomanufacturing facilities in a matter of days or weeks (McDonald and Holtz, 2020). As an exercise to practically test the repurposing requirements, we describe here the partial adaptation of our research laboratory and greenhouse facilities to the production of SARS-CoV-2-related antigens and antibodies using *N. benthamiana* agroinfiltration as manufacturing platform.

In Figure 6 we show a chronogram of the activities undertaken by our team towards the production of SARS-CoV-2 antigens and antibodies, from the initial selection of the nucleotide sequences of the genes-of-interest to the production of one litre of plant apoplastic fluid of recombinant sybody17 and nanobody72. In our hands, the whole process took a total of nine weeks with non-exclusive personnel dedication and partially restricted access to our facilities. The process can be divided in three periods: the first step (DESIGN), taking approximately ten days, was dedicated to construct design and gene synthesis. It was pivotal in this step to have open access to viral and antibody sequences deposited in pre-print repositories. Particularly remarkable was the openness of academic labs that immediately released primary sequence information of partially characterized anti-SARS-CoV-2 monoclonal antibodies, an exercise that should serve as an example in the future. Due to our limited testing capacity, the number of parallel designs per product was maintained relatively low, and several design decisions (e.g. codon optimization, purification tags) were taken based in a best-guess approach. Ideally, proper crisis preparedness should involve a centralized automated equipment such as a biofoundry (Hillson et al., 2019), with which the design space could be extended dramatically without causing delay. The second phase (BUILD) was dedicated to cloning and construct building and lasted less than three weeks. Our lab counts with adapted plasmids and cloning procedures from previous projects (Sarrion-Perdigones et al., 2011; Vazquez-Vilar et al., 2017), therefore no significant time lag occurred in this step. Importantly, this period also involved seeding a new plant batch at the scale required for pilot production in week seven (112 plants distributed in 16 hexagonal modules in this case). In a third phase (TEST), starting on week 5, all constructs were infiltrated at a small scale (three replicate leaves each), shortly incubated (5 or 7 days) and then tested functionally in parallel analyses. This small-scale assay took two additional weeks, summing a total of approximately 50 days for the complete process. The Synthetic Biology-inspired Design-Build-Test (DBT) process described above is conceived as an iterative one, so that new DBT cycles can be run fuelled by the conclusion of previous cycles to generate new optimized versions of the product. Based on this experience, we estimate that the whole DBT cycle could be shortened to 30 days or less by optimising the pipeline (e.g. introducing centralized, automatized design and build phases), and by improving preparation and anticipation in the facilities (Fig 6). For instance, note that moving from step 2 to step 3 without delay requires a small batch of plants be always maintained in the facility, as it was in our case to supply our research requirements. This only involves transplanting 10-15 seedlings every three weeks, and then disposing of them every other three weeks once they start flowering. If a continuous plant supply is not maintained, a minimum of three extra weeks needs to be considered to have plants ready for the first TEST iteration.

**Figure 6.**
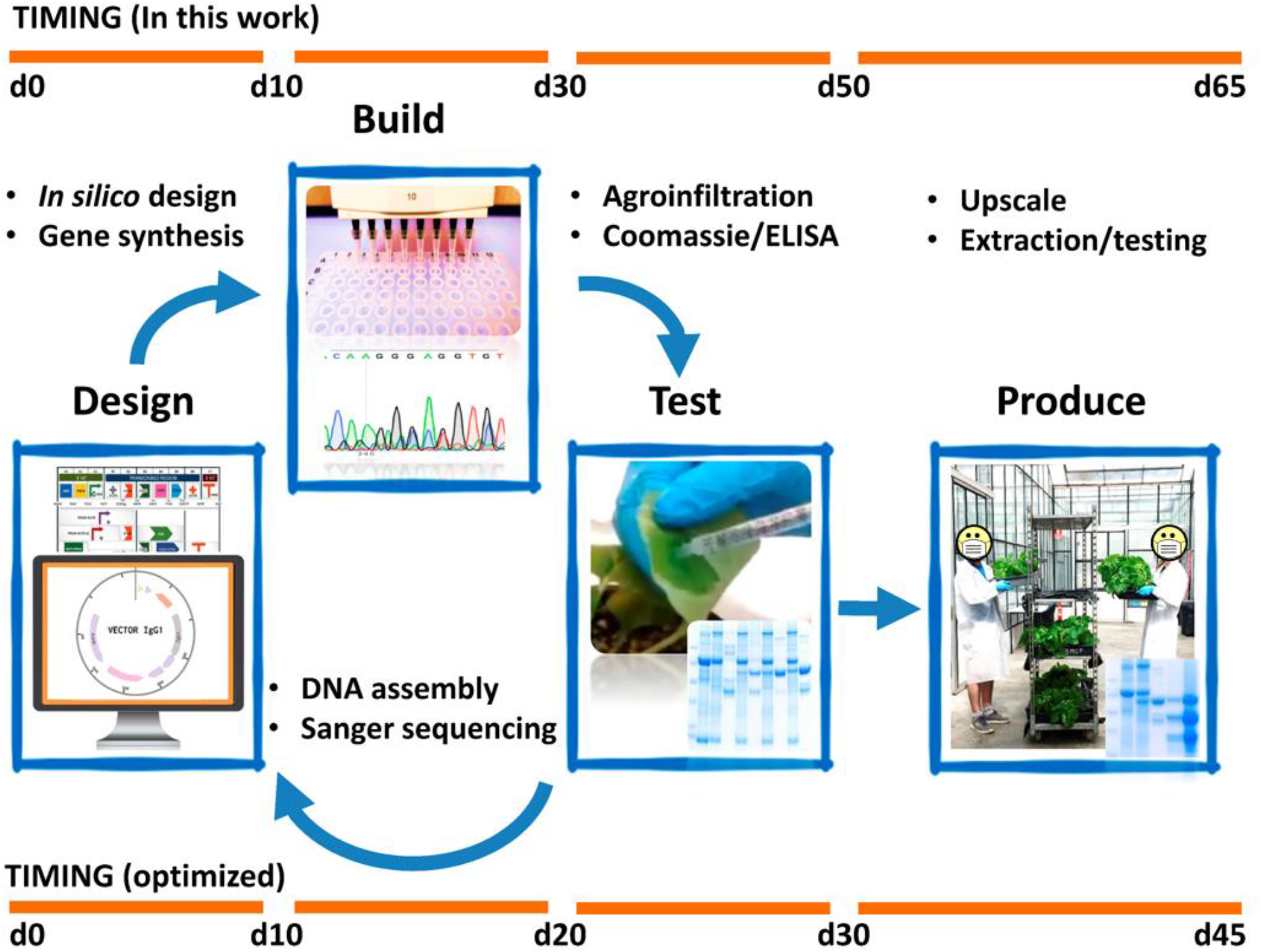
Schematic view of the Design-Build-Test and Production pipelines employed in this work. The upper timeline represents the actual timespan of the experiments. Note that the time points represent approximately the days required to produce and initially characterize the designated products; notwithstanding, some of the results shown in previous figures correspond to extended analysis obtained at a later stage, during the preparation of this manuscript. The lower timeline describes the estimated minimal timespan that would result by introducing some of the optimizations described in the text.

Whereas the first version of products shown here lack iterative optimization, it would serve eventually to respond to the most urgent demands. In our case, as the results of the first DBT process arose, the best performing version (V1.0) of two of the products were taken to PRODUCTION phase. In the exercise shown here, the upscaling was relatively small (112 plants, approximately 2.5 Kg FW). Post-agroinfiltration incubation time was extended to 14 days to maximize yields. In the meantime, optimization of the purification/extraction methods were undertaken at small scale, so that the new knowledge acquired could be applied in the batch purification of the pilot experiment. In a crisis-scenario, and given the modularity of the proposed scheme, several medium-size production modules can be replicated in a farming facility, and reproduced in several farms, allowing easy scalability. Successive iterations with small scale agroinfiltration could be an effective way to maximize yields and reduce development times by comparing different small-scale strategies. It should be mentioned that the basic apoplast-based downstream processing proposed here could only be used, with the necessary adaptations, in a limited number of crisis-related applications, mainly related with detection and diagnosis. Other uses, certainly therapeutic ones, would involve additional regulatory considerations including GMP downstream facilities, which are beyond the scope of this exercise.

As a result of this experience, several improvements can be envisioned. We employed the MagnICON® vector system with few adaptations for all the attempted proteins. Although MagnICON® produces maximum yields for many products, some proteins, particularly viral antigens may express better with other (e.g. non-viral) systems. In our experiments, antigens showed rather low expression levels despite optimization attempts using codon optimization and different localization signals. In adapting to an emergency, it would be advisable to perform initial expression tests using different production platforms also involving non-replicative methods (Sainsbury and Lomonossoff, 2008) or DNA viruses (Yamamoto et al., 2018; Zhang and Mason, 2006), and to incorporate them to the initial optimization test. As mentioned, this could be done in a centralized manner, later distributing expression clones to several repurposed production facilities. In contrast to antigens, recombinant single-chain antibodies showed in general higher and more uniform expression levels, as could be expected from their more similar structure. We chose to adapt full human IgGs to a scFv-IgG1 format to facilitate cloning and expression procedures, since it has been earlier described in plants that single chain formats reproduce the binding activities of the original full-size antibodies from which they derive. This format also facilitates comparisons with VHH antibodies, also produced as IgG1 fusions.

The plant-made SARS-CoV-2 products described here have several potential applications in the diagnosis area. Both RBD and N proteins can be used as reagents for serological assays (Amanat et al., 2020; Liu et al., 2020), although further yield optimizations should be required. For those assays where antigen glycosylation is an important factor, glycoengineered plants (Strasser et al., 2008) can provide a competitive alternative to mammalian cells cultures (O’Flaherty et al., 2020). Regarding antibodies, they can serve as internal references for the quantification of serological responses. With small modifications, the same antibodies can be adapted for sandwich ELISA and employed in the detection and quantification of viral particles, a better proxy for infectiveness than RNA. We also show here that apoplastic fluid is an inexpensive antibody preparation suitable for certain applications that require low-cost preparations, e.g. the concentration of the virus from environmental samples. As shown here, the protein complexity in the apoplast is greatly reduced, therefore the apoplast could be regarded as a plant-equivalent of hybridoma supernatant or ascited fluid, although at much lower cost. Unfortunately, apoplastic preparations are prone to partial antibody degradation, probably due to endogenous proteases, however this can be minimized using extraction buffers with appropriate protease inhibitors, as it was shown for nanobody72.

The current pandemic crisis has evidenced the power of new antibody selection procedures, either based on single-cell selection from human peripheral blood mononuclear cells, in the case of full-size antibodies, or based on ultra-high throughput selection of synthetic libraries (sybodies) in the case of camelid-derived nanobodies (Zimmermann et al., 2018; Walter et al., 2020). Large collections of anti-SARS-CoV-2, potentially neutralizing antibody sequences were made available to the scientific community in a question of weeks rather than months. It does not go unnoticed that the combination of rapid antibody selection procedures with fast, modular and scalable plant expression also has implications in the therapeutic arena as an ideal system for passive immunization. Intravenous polyclonal immunoglobulins (IVIG) from recovered patients have been shown a very effective CoVid-19 treatment in several studies (Montelongo-Jauregui et al., 2020, and references herein) however the limited availability of patient sera hampers its application in practice. Interestingly, we showed in a recent work that large recombinant polyclonal antibody cocktails (pluribodies), mimicking a mammalian immune response can be produced in *N. benthamiana* with high batch-to-batch reproducibility (Julve Parreño et al., 2018). Passive immunization with recombinant antibody cocktails resembles a natural response more than a monoclonal therapy, requires shorter developmental times and is probably more robust against the development of resistances.

In conclusion, based on the results of the exercise described here, we propose the repurposing of indoors farms into plant-based biomanufacturing facilities as a viable option to respond to local and global shortages of bioproducts such as diagnostics and therapeutic reagents in times of crisis.

## AUTHOR CONTRIBUTIONS

All authors designed and performed the experiments, and analyzed the data. D.O. wrote this manuscript. All authors revised and edited the written manuscript.

## ACKNOWLEDGEMENTS

The authors want to thank to the staff of the IBMCP and the Polytechnic University of Valencia who help us to access safely to our laboratory and greenhouses during the CoVid-19 lockdown in Spain, and specially Eugenio Grau for his readiness to help us with Sanger sequencing during that period. We are grateful to Prof. Steinkellner and Strasser for providing glycoengineered plant lines and to Prof. Gleba for sharing Magnifection plasmids.

## REFERENCES

Amanat, F., Stadlbauer, D., Strohmeier, S., Nguyen, T. H. O., Chromikova, V., McMahon, M., et al. (2020). A serological assay to detect SARS-CoV-2 seroconversion in humans. Nat Med 26, 1033–1036. doi:10.1038/s41591-020-0913-5.

Armbruster, D. A., and Pry, T. (2008). Limit of Blank, Limit of Detection and Limit of Quantitation. Clin. Biochem. Rev 29, S49–S52.

Buyel, J. F. (2019). Plant Molecular Farming – Integration and Exploitation of Side Streams to Achieve Sustainable Biomanufacturing. Front. Plant Sci. 9, 1893. doi:10.3389/fpls.2018.01893.

Capell, T., Twyman, R. M., Armario-Najera, V., Ma, J. K.-C., Schillberg, S., and Christou, P. (2020). Potential Applications of Plant Biotechnology against SARS-CoV-2. Trends Plant Sci. doi:10.1016/j.tplants.2020.04.009.

Dobon, A. (2019). Transient Gene Expression Seeds Plant-Based Bioproduction Systems: Leaf Expression Systems’ Hypertrans technology promises low costs and high yields. Genetic Engineering & Biotechnology News 39, 54–56. doi:10.1089/gen.39.03.14.

Fernandez-del-Carmen, A., Juárez, P., Presa, S., Granell, A., and Orzáez, D. (2013). Recombinant jacalin-like plant lectins are produced at high levels in Nicotiana benthamiana and retain agglutination activity and sugar specificity. Journal of Biotechnology 163, 391–400. doi:10.1016/j.jbiotec.2012.11.017.

Giritch, A., Marillonnet, S., Engler, C., van Eldik, G., Botterman, J., Klimyuk, V., et al. (2006). Rapid high-yield expression of full-size IgG antibodies in plants coinfected with noncompeting viral vectors. Proceedings of the National Academy of Sciences 103, 14701–14706. doi:10.1073/pnas.0606631103.

Gleba, Y., Marillonnet, S., and Klimyuk, V. (2004). Engineering viral expression vectors for plants: the ‘full virus’ and the ‘deconstructed virus’ strategies. Current Opinion in Plant Biology 7, 182–188. doi:10.1016/j.pbi.2004.01.003.

Hillson, N., Caddick, M., Cai, Y., Carrasco, J. A., Chang, M. W., Curach, N. C., et al. (2019). Building a global alliance of biofoundries. Nat Commun 10, 2040. doi:10.1038/s41467-019-10079-2.

Julve, J. M., Gandía, A., Fernández-del-Carmen, A., Sarrion-Perdigones, A., Castelijns, B., Granell, A., et al. (2013). A coat-independent superinfection exclusion rapidly imposed in Nicotiana benthamiana cells by tobacco mosaic virus is not prevented by depletion of the movement protein. Plant Mol Biol 81, 553–564. doi:10.1007/s11103-013-0028-1.

Julve Parreño, J. M., Huet, E., Fernández-del-Carmen, A., Segura, A., Venturi, M., Gandía, A., et al. (2018). A synthetic biology approach for consistent production of plant-made recombinant polyclonal antibodies against snake venom toxins. Plant Biotechnol J 16, 727–736. doi:10.1111/pbi.12823.

Liu, W., Liu, L., Kou, G., Zheng, Y., Ding, Y., Ni, W., et al. (2020). Evaluation of Nucleocapsid and Spike Protein-Based Enzyme-Linked Immunosorbent Assays for Detecting Antibodies against SARS-CoV-2. J Clin Microbiol 58, e00461–20, /jcm/58/6/JCM.00461-20.atom. doi:10.1128/JCM.00461-20.

Marillonnet, S., Thoeringer, C., Kandzia, R., Klimyuk, V., and Gleba, Y. (2005). Systemic Agrobacterium tumefaciens–mediated transfection of viral replicons for efficient transient expression in plants. Nat Biotechnol 23, 718–723. doi:10.1038/nbt1094.

McDonald, K. A., and Holtz, R. B. (2020). From Farm to Finger Prick—A Perspective on How Plants Can Help in the Fight Against COVID-19. Front. Bioeng. Biotechnol. 8, 782. doi:10.3389/fbioe.2020.00782.

Montelongo-Jauregui, D., Vila, T., Sultan, A. S., and Jabra-Rizk, M. A. (2020). Convalescent serum therapy for COVID-19: A 19th century remedy for a 21st century disease. PLOS Pathogens 16, e1008735. doi:10.1371/journal.ppat.1008735.

Moon, K.-B., Park, J.-S., Park, Y.-I., Song, I.-J., Lee, H.-J., Cho, H. S., et al. (2019). Development of Systems for the Production of Plant-Derived Biopharmaceuticals. Plants 9, 30. doi:10.3390/plants9010030.

O’Flaherty, R., Bergin, A., Flampouri, E., Mota, L. M., Obaidi, I., Quigley, A., et al. (2020). Mammalian cell culture for production of recombinant proteins: A review of the critical steps in their biomanufacturing. Biotechnology Advances 43, 107552. doi:10.1016/j.biotechadv.2020.107552.

Olinger, G. G., Pettitt, J., Kim, D., Working, C., Bohorov, O., Bratcher, B., et al. (2012). Delayed treatment of Ebola virus infection with plant-derived monoclonal antibodies provides protection in rhesus macaques. Proceedings of the National Academy of Sciences 109, 18030–18035. doi:10.1073/pnas.1213709109.

Sainsbury, F., and Lomonossoff, G. P. (2008). Extremely High-Level and Rapid Transient Protein Production in Plants without the Use of Viral Replication. Plant Physiol. 148, 1212–1218. doi:10.1104/pp.108.126284.

Sainsbury, F., Thuenemann, E. C., and Lomonossoff, G. P. (2009). pEAQ: versatile expression vectors for easy and quick transient expression of heterologous proteins in plants. Plant Biotechnology Journal 7, 682–693. doi:10.1111/j.1467-7652.2009.00434.x.

Sarrion-Perdigones, A., Falconi, E. E., Zandalinas, S. I., Juárez, P., Fernández-del-Carmen, A., Granell, A., et al. (2011). GoldenBraid: An Iterative Cloning System for Standardized Assembly of Reusable Genetic Modules. PLoS ONE 6, e21622. doi:10.1371/journal.pone.0021622.

Sarrion-Perdigones, A., Vazquez-Vilar, M., Palaci, J., Castelijns, B., Forment, J., Ziarsolo, P., et al. (2013). GoldenBraid 2.0: A Comprehensive DNA Assembly Framework for Plant Synthetic Biology. PLANT PHYSIOLOGY 162, 1618–1631. doi:10.1104/pp.113.217661.

Strasser, R., Stadlmann, J., Schähs, M., Stiegler, G., Quendler, H., Mach, L., et al. (2008). Generation of glyco-engineered Nicotiana benthamiana for the production of monoclonal antibodies with a homogeneous human-like N-glycan structure: XylT and FucT down-regulation in N. benthamiana. Plant Biotechnology Journal 6, 392–402. doi:10.1111/j.1467-7652.2008.00330.x.

Tian, X., Li, C., Huang, A., Xia, S., Lu, S., Shi, Z., et al. (2020). Potent binding of 2019 novel coronavirus spike protein by a SARS coronavirus-specific human monoclonal antibody. Emerging Microbes & Infections 9, 382–385. doi:10.1080/22221751.2020.1729069.

van den Brink, E. N., ter Meulen, J., Cox, F., Jongeneelen, M. A. C., Thijsse, A., Throsby, M., et al. (2005). Molecular and Biological Characterization of Human Monoclonal Antibodies Binding to the Spike and Nucleocapsid Proteins of Severe Acute Respiratory Syndrome Coronavirus. JVI 79, 1635–1644. doi:10.1128/JVI.79.3.1635-1644.2005.

Vazquez-Vilar, M., Quijano-Rubio, A., Fernandez-del-Carmen, A., Sarrion-Perdigones, A., Ochoa-Fernandez, R., Ziarsolo, P., et al. (2017). GB3.0: a platform for plant bio-design that connects functional DNA elements with associated biological data. Nucleic Acids Res, gkw1326. doi:10.1093/nar/gkw1326.

Walter, J. D., Hutter, C. A. J., Zimmermann, I., Wyss, M., Egloff, P., Sorgenfrei, M., et al. (2020). Sybodies targeting the SARS-CoV-2 receptor-binding domain. Biochemistry doi:10.1101/2020.04.16.045419.

Wirz, H., Sauer-Budge, A. F., Briggs, J., Sharpe, A., Shu, S., and Sharon, A. (2012). Automated Production of Plant-Based Vaccines and Pharmaceuticals. J Lab Autom. 17, 449–457. doi:10.1177/2211068212460037.

Wrapp, D., De Vlieger, D., Corbett, K. S., Torres, G. M., Wang, N., Van Breedam, W., et al. (2020). Structural Basis for Potent Neutralization of Betacoronaviruses by Single-Domain Camelid Antibodies. Cell 181, 1004–1015.e15. doi:10.1016/j.cell.2020.04.031.

Yamamoto, T., Hoshikawa, K., Ezura, K., Okazawa, R., Fujita, S., Takaoka, M., et al. (2018). Improvement of the transient expression system for production of recombinant proteins in plants. Scientific Reports 8, 4755. doi:10.1038/s41598-018-23024-y.

Zhang, X., and Mason, H. (2006). Bean Yellow Dwarf Virus replicons for high-level transgene expression in transgenic plants and cell cultures. Biotechnol. Bioeng. 93, 271–279. doi:10.1002/bit.20695.

Zimmermann, I., Egloff, P., Hutter, C. A., Arnold, F. M., Stohler, P., Bocquet, N., et al. (2018). Synthetic single domain antibodies for the conformational trapping of membrane proteins. eLife 7, e34317. doi:10.7554/eLife.34317.

Zrein, M., De Marcillac, G., and Van Regenmortel, M. H. V. (1986). Quantitation of rheumatoid factors by enzyme immunoassay using biotinylated human IgG. Journal of Immunological Methods 87, 229–237. doi:10.1016/0022-1759(86)90536-3.

