## Supplementary Table 1 for "Pilot production of SARS-CoV-2 related proteins in plants: a proof of concept for rapid repurposing of indoors farms into biomanufacturing facilities"

### *Supplementary Material*

**Supplementary Table 1.** List of Level 0 plasmids generated during this work. All sequences can be searched at https://gbcloning.upv.es/search/features/ using the GB ID.

| Name | GB ID |
|---|---|
| **pUPD2 CR3009** | GB3576 |
| **pUPD2 CR3018** | GB3575 |
| **pUPD2 CR3022** | GB3577 |
| **pUPD2 Sybody3** | GB3402 |
| **pUPD2 Sybody17** | GB3403 |
| **pUPD2 Nanobody72** | GB3404 |
| **pUPD2 natRBD:His** | GB3377 |
| **pUPD2 bcoRBD:His** | GB3382 |
| **pUPD2 His:natRBD:KDEL** | GB3441 |
| **pUPD2 His:bcoRBD:KDEL** | GB3442 |
| **pUPD2 His:bcoN** | GB3447 |
